# The spread of *Aedes albopictus* in the islands of São Tomé and Príncipe

**DOI:** 10.1101/2023.09.28.559833

**Authors:** Jonathan A. Rader, Antonio Serrato-Capuchina, Tayte Anspach, Daniel R. Matute

## Abstract

The mosquito *Aedes albopictus* is a vector species of Dengue, yellow fever, and Zika among other diseases. The species originated in Southeast Asia and has spread widely and rapidly in the last century. The species has been reported in localities from the Gulf of Guinea since the early 2000s, but systematic sampling has been scant. We sampled *Ae. albopictus* between 2013 and 2023 across the altitudinal gradient in São Tomé and found that the species was present in all sampled years at altitudes up to 680 meters. We also find some evidence of increases in proportional representation compared to *Ae. aegypti* over time. We report the presence of the species in Príncipe for the first time, suggesting that the range of *Ae. albopictus* is larger than previously thought. Finally, we use bioclimatic niche modeling to infer the potential range of *Ae. albopictus* and infer that the species has the potential to spread across a large portion of São Tomé and Príncipe. Our results suggest that *Ae. albopictus* has established itself as a resident species of the islands of the Gulf of Guinea and should be incorporated into the list of potential vectors that need to be surveyed and controlled.

## INTRODUCTION

Mosquitoes of the genus *Aedes* (Diptera: Culicidae) are some of the most important disease vector species. The genus is endemic to the Old World and encompasses over 700 species (Das *et al*. 2018), of which at least 12 have been determined to be vectors of human illnesses such as dengue and yellow fever (Mattingly 1967; Jupp and Kemp 2002; Molaei *et al*. 2008; Benelli and Romano 2017). Dengue alone is estimated to cause over 390 million infections per year, of which 96 million have clinical manifestations (Bhatt *et al*. 2013). Over half of the world’s population (3.9 billion people) are at risk of infection with dengue and the risk of contracting the disease has increased 30-fold worldwide in the last 50 years (Organization and Asia 2011). Each year at least 50,000 patients die because of dengue (Wilder-Smith *et al*. 2010) and at least 30,000 because of yellow fever (Barnett 2007). These numbers are, in all likelihood, underestimates of the actual *Aedes*-borne disease burden (Rigau-Pérez 2006; Barnett 2007; Allwinn *et al*. 2008; Adalja *et al*. 2012; Beaumier *et al*. 2014). *Aedes* mosquitoes also transmit other pathogens that cause severe diseases including four more arboviruses (Mayaro, West Nile fever, eastern equine encephalitis, and Zika) and in some cases filarial diseases (Wharton *et al*. 1963). As climate change increases temperatures worldwide, the range of *Aedes* continues to extend across each continent (Kraemer *et al*. 2019a). Quantifying novel invasions of *Aedes* is vital to mitigating risk towards human health worldwide.

Two species are the most notable vectors of disease in the group, *Aedes aegypti* and *Aedes albopictus*. *Aedes aegypti* is endemic to Northwest Africa, and it has been speculated that the species expanded to the Americas during the TransAtlantic slave trade (Brown *et al*. 2014; Pless *et al*. 2022). Genome sequencing also suggests that specialization to human habitats arose as an adaptation to hot and dry seasons in the West African Sahel; the water store in human settlements acted as an oviposition habitat for *Ae. aegypti* (Rose *et al*. 2023). *Aedes albopictus* is native to Southeast Asia but has expanded its range considerably over the past few decades and has been hypothesized to be driven by human activities (Bonizzoni *et al*. 2013; Goubert *et al*. 2016; Kotsakiozi *et al*. 2017). Human migration within the Indo-Malayan Peninsula and the Indian Ocean islands, including Madagascar, favored the early spread (i.e., potentially as early as 1904, (Fontenille and Rodhain 1989; Longbottom *et al*. 2023)) of *Ae. albopictus* into new ranges with similar biogeographic conditions. The spread was exacerbated by the explosion of intercontinental trade in the 20th century and, in particular, the shipment of tires among countries seems to have been responsible for the introduction of *Ae*. *albopictus* into the USA circa 1986 (Sprenger and Wuithiranyagool 1986; Moore and Mitchell 1997). Systematic sampling of the presence of *Aedes* species has indicated that the two main vector species of *Aedes* are rapidly increasing their geographic range (Ryan *et al*. 2019; Kraemer *et al*. 2019a; Longbottom *et al*. 2023).

Biological invasions are important to understand how adaptation to novel environments occurs, but are also crucial to understanding what delimits ranges of disease vectors. Biological invasions are characterized by rapid range expansion following introduction. *Ae. aegypti* is almost globally distributed following introductions beginning in the last few centuries (Powell and Tabachnick 2013; Brown *et al*. 2014; Pless *et al*. 2022). *Aedes albopictus* is characterized as one of the most contemporarily invasive species as it has it expanded its range rapidly in the last few decades (reviewed in (Benedict *et al*. 2007; Longbottom *et al*. 2023)); its invasive nature combined with its vector competency (Mitchell 1995; Jupp and Kemp 2002; Gloria-Soria *et al*. 2020) makes it a concern for many countries. Even though this species was initially considered a secondary vector of yellow fever, and has been suggested as a strong heterospecific competitor that might assist in the eradication of *Ae. aegypti* (Juliano 2010), the last years have revealed that *Ae. albopictus* can serve as a successful vector of Zika and Chikunguya (Jupp and Kemp 2002; Garcia-Luna *et al*. 2018; McKenzie *et al*. 2019). A precise description of the potential suitable habitats of each of these two species is sorely needed to identify the factors that can constrain the spread of these vectors.

Environmental niche modeling suggests these vectors might expand into large swaths of the globe (Kutsuna *et al*. 2015; Kamal *et al*. 2018) and indeed in the last 25 years, the rate of range expansion of *Ae. aegypti* in places like North America has reached speeds of 250 km/year (Kraemer *et al*. 2019b). The rate of *Ae. albopictus* range expansion is also high and currently ranges between 60 km and ∼150 km per year depending on the geographic location (Kraemer *et al*. 2019b). As *Aedes* species have expanded their ranges, the number of people exposed to the diseases they vector has increased drastically in the last 10 years (Bouri *et al*. 2012; Adalja *et al*. 2012; Beaumier *et al*. 2014; Rey 2014; Stanaway *et al*. 2016; Ryan *et al*. 2019). This range expansion has coincided with changes in the mosquitos’ selective environment, including anthropogenic selective pressures such as insecticide exposure (Armstrong *et al*. 2017), availability of substrates (Abreu *et al*. 2015), and host density (Rodrigues *et al*. 2015). The range of these species is expected to keep growing in the future (Mogi *et al*. 2017; Kraemer *et al*. 2019b). To understand the distribution of the pathogens vectored by mosquitoes, systematic local-scale data collection is sorely needed. In particular, the relatively closed environments such as oceanic islands provide “natural experiments” that can illuminate how novel species move across isolating geographic barriers, and how genetic variability accrues over time in the newly-founded population. Moreover, instances of recent invasions can reveal the dynamics of how vector species become established.

In this piece, we followed up on the report that *Ae. albopictus* is present in São Tomé island (Reis *et al*. 2017) and study the presence of *Aedes albopictus* across multiple locations of São Tome, including at high altitude sites (∼1,400 meters above the sea level). We also report the first observation of this vector on the island of Príncipe. We used occurrence data to infer the current ranges of the two species and determine whether they are likely to increase their ranges as global temperatures rise. These observations add to the body of work that indicates that *Ae*. *albopictus* has colonized the islands of the Gulf of Guinea (Toto *et al*. 2003) and poses an additional challenge on vector surveillance on the islands of the Cameroon Volcanic line in the Gulf of Guinea.

## METHODS

### Collections

Our first collections of *Aedes* in São Tomé were carried out in Montecafe, a mid-elevation town in São Tomé, in puddles of standing water (Figure 1) in the year 2013. We collected larvae using Mosquito dippers powdered with stainless steel (John W. Hock Company, Gainesville, FL) and aspirated adults with mouth aspirators and CDC-type light traps (2836BQX, BioQuip; Rancho Domingo, CA) connected to a 6V as a power source. We repeated the same sampling in 2023. We used a yeast-water paste as a source of CO_2_. Traps were emptied every 48 hours, when specimens were collected for identification, and the battery was replaced. Traps were kept in place for a total of 10 days but were sampled daily to collect specimens. Yeast paste around the traps was replaced daily. The sampling in 2013 took place in June-July and the sampling in 2023 took place in January.

**FIGURE 1.**
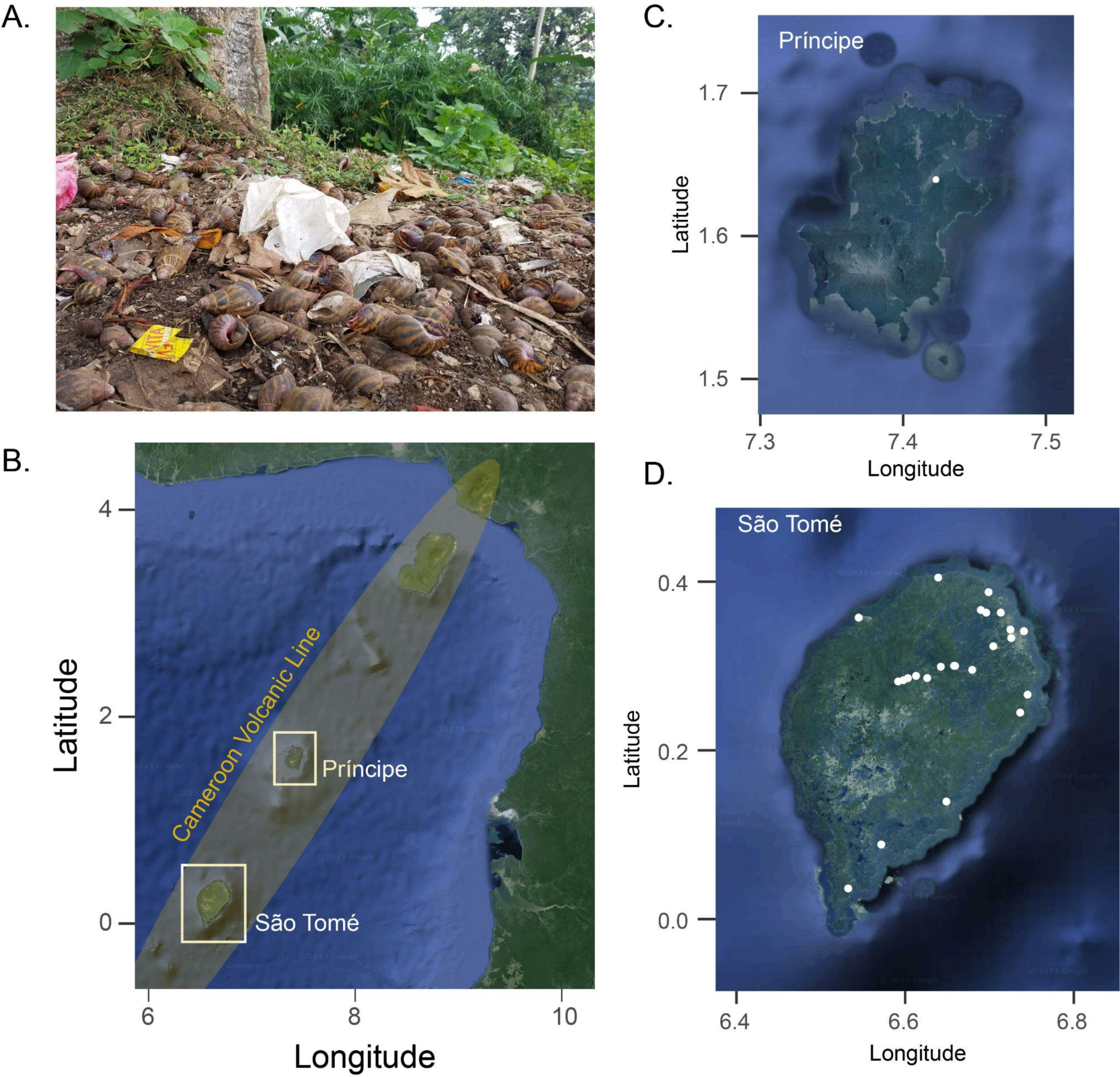
Mosquito collection scheme. A. Example of a location that yielded *Ae. albopictus* in the midlands of São Tomé. B. Map of the Gulf of Guinea, depicting the islands of the Carmeroon Volcanic Line and highlighting São Tomé and Príncipe. C. Map showing the location of the traps on Príncipe and on São Tomé (D.).

### Species identification: morphology

Larvae were inspected for morphological traits which differentiate between *Ae. aegypti* and *Ae. albopictus*. All larvae were inspected using a non-destructive procedure. We differentiated between *Ae. albopictus* and *Ae. aegypti* larvae using two diagnostic traits. In *Ae. aegypti*, the comb scale on the terminal segment has a single row of pitchfork-shaped scales; larvae also have strong black hooks in the side of the thorax. In *Ae. albopictus*, the comb scale in the terminal segment is also in one row but the scales are thorn-shaped and the hooks on the side of the thorax are small or absent (Lee 1998). Pupae were differentiated by the shape and length of the terminal setae in the paddle. While *Ae. aegypti* has short and simple terminal setae at the edge of the paddle, *Ae. albopictus* has long hairs (Penn 1949). Larvae and pupae were kept in glass vials until they hatched and were inspected for adult traits. In particular we studied traits that differentiate between these two species. In *Ae. albopictus* the scutum has one silvery-white stripe down its middle, the thorax has patches of silver scales and a black clipeus. In *Ae. aegypti*, the scutum has a lyre-shaped pattern of white scales, a clipeus with white scales. We collected no samples of other species that have been reported in the island (*Aedes nigricephalus, Aedes circumluteolus, and Aedes metallicus,* (Loiseau *et al*. 2022)) or other species that are common in the continent (i.e., *Ae. africanus*).

### Altitudinal gradients

We used a linear model in which abundance was the response, and altitude and species were the factors of the model. The model took the form:

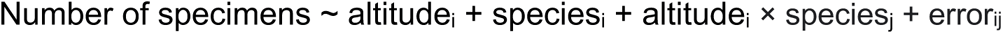

To assess significance of the effects we used the R function *anova* (library *stats*, (R Core Team 2018).

Finally, to determine if the proportional representation of *Ae. albopictus* had changed between 2013 and 2023, we calculated the proportion of *Ae. albopictus* at 13 locations for larvae and 15 locations for adults. Other high altitude locations lacked sufficient specimens for a comparison. To determine if there had been an increase in the proportion of *Ae. albopictus*, we used a one tailed t-test comparing the change to 0 (R function *t.test*, library *stats*, (R Core Team 2018)).

### Species identification: DNA barcoding

To confirm the identification using morphological traits, we used a mtDNA marker, cytochrome oxidase I. Previous studies have shown that COI-based DNA barcoding is able to differentiate between species of *Aedes*, and other genera of mosquitoes (Chan *et al*. 2014). We used the two following primers ‘-AAAAAGATGTATTTAAATTTCGGTCTG-3’ and 5’-TGTAATTGTTACTGCTCATGCTTTT-3’. We extracted DNA from single mosquitoes using the Qiagen DNeasy Blood and Tissue Kit (Qiagen Inc., Valencia, CA). We then amplified ∼25 ng of total DNA using PCR and conditions previously reported (Chan *et al*. 2014). Briefly the amplification cycles were as follows: 94°C for 1 min followed by (34 cycles at 94°C for 30 s, annealing temperature: 52°C for 30 s, 72°C for 1 min 30s), and a single final step at 72°C for 7 min. All amplifications were performed using a 2720 Thermal Cycler from Applied Biosystems (Foster City, CA). PCR products were sent for sequencing to Eton Biosciences (Research Triangle Park, NC) using the same PCR primers. Sequencing of PCR fragments was performed using the cyclic reaction termination method (Sanger sequencing) using the BigDye Terminator Cycle Sequencing Kits (Applied Biosystems, Foster City, CA, USA). These sequencing yielded amplicons for 177 individuals from five locations in São Tomé (*n* = 155) and Príncipe (*n* = 12). To minimize false polymorphism, we sequenced both DNA strands, aligned them, and examined them with 4Peaks 1.7.1 (http://nucleobytes.com/index.php/4peaks). The sequences of novel unique haplotypes were deposited in Genbank (TBD).

Next, we downloaded previously COI data for other species of *Aedes* (accession numbers are listed in Table S1), and aligned the 47 DNA sequences (six from this study and 41 from previous studies, including one for the outgroup, *Ae. aegypti*) using ClustalX (Larkin et al., 2007). From the PCR fragments we amplified, we only used unique haplotypes. We generated a COI-base gene genealogy and assessed the morphological identification of the samples in São Tomé and Principe samples. We used IQ-TREE (Nguyen *et al*. 2015) to generate the tree using the following using-m MFP option (ModelFinder, (Kalyaanamoorthy *et al*. 2017)) to automatically select the best-fitting sequence evolution model. To measure individual tree branch support, we ran 1,000 ultrafast bootstrap replicates and followed them with a Shimodaira–Hasegawa-like approximate likelihood ratio test (SH-aLRT, (Minh *et al*. 2013)).

### Thermal niche and distribution

São Tomé and Príncipe are both small islands (855 km^2^ and 109 km^2^, respectively) but encompass a large variety of ecological niches. We examined the potential thermal niche of *Ae. albopictus* on the islands of São Tomé and Príncipe. Augmented with climate information sourced from the WorldClim database (Fick and Hijmans 2017; Cerasoli *et al*. 2022), we sought to create a comprehensive portrayal of their preferred habitats. Leveraging the raster R library, as detailed by Hijmans et al. 2019, we focused on bioclim variables at a resolution of 30 arc seconds. We constructed a Maximum Entropy model (MaxEnt; (Phillips *et al*. 2004, 2006) using global occurrence data for *Ae. albopictus*, excluding the samples we report here for the islands of São Tomé and Príncipe. Our model is based on 79,247 observations from the Global Biodiversity Information Facility (GBIF, (Telenius 2011; Luo *et al*. 2021)) from 1980 to the present (see Figure S1). We included 17 of the 19 bioclimatic variables from WorldClim (Fick and Hijmans 2017; Cerasoli *et al*. 2022), omitting composite variables to avoid multicollinearity (see Table S2). We used the ‘*maxent*’ command in the R *dismo* package (Hijmans *et al*. 2017) to build a global MaxEnt model with 30 arcsecond resolution, which we then cropped to São Tomé and Príncipe to predict the proportion of the islands that may provide suitable habitat for *Ae. albopictus*.

We used the MaxEnt model to describe the bioclimatic niche availability of *Ae. albopictus* on the islands of São Tomé and Príncipe. To approximate the global bioclimatic niche for *Ae. albopictus*, we calculated the middle 95% quantile ranges for the five Worldclim variables with the greatest contributions to our model (see Figure 5 and Table S2). We discarded the upper and lower 2.5% quantiles. To assess the availability of suitable bioclimatic zones on the islands, we cropped the rasters for the MaxEnt predictors to each of the islands, extracted the range of values for each of the five variables, and assessed the overlap of the island values with the global fundamental niche. Finally, we overlaid our samples onto the MaxEnt output for each variable and extracted the bioclimatic niche breadth currently encompassed by the known range of *Ae. albopictus* on the islands.

## RESULTS

First, we collected larvae and adults from standing water and using aspirators in the lowlands and the midlands of the island of São Tomé in 2013. High elevations (over 700 meters above the sea level) yielded no vector collections. We found that in the island of São Tomé both, *Ae. aegypti* and *Ae. albopictus*, were present as adults at altitudes of 891 meters in 2023. Our collections confirm the observation that *Ae. albopictus* has been present in São Tomé island since at least 2013, remained present in 2016 (Reis *et al*. 2016), and was still present in 2023 (Figure 2). *Aedes albopictus* was present at all but the five highest elevation sample sites (high elevation sites ranged from 1385.5 to 2825.1 m), but were present up to approximately 1330 m.

**FIGURE 2.**
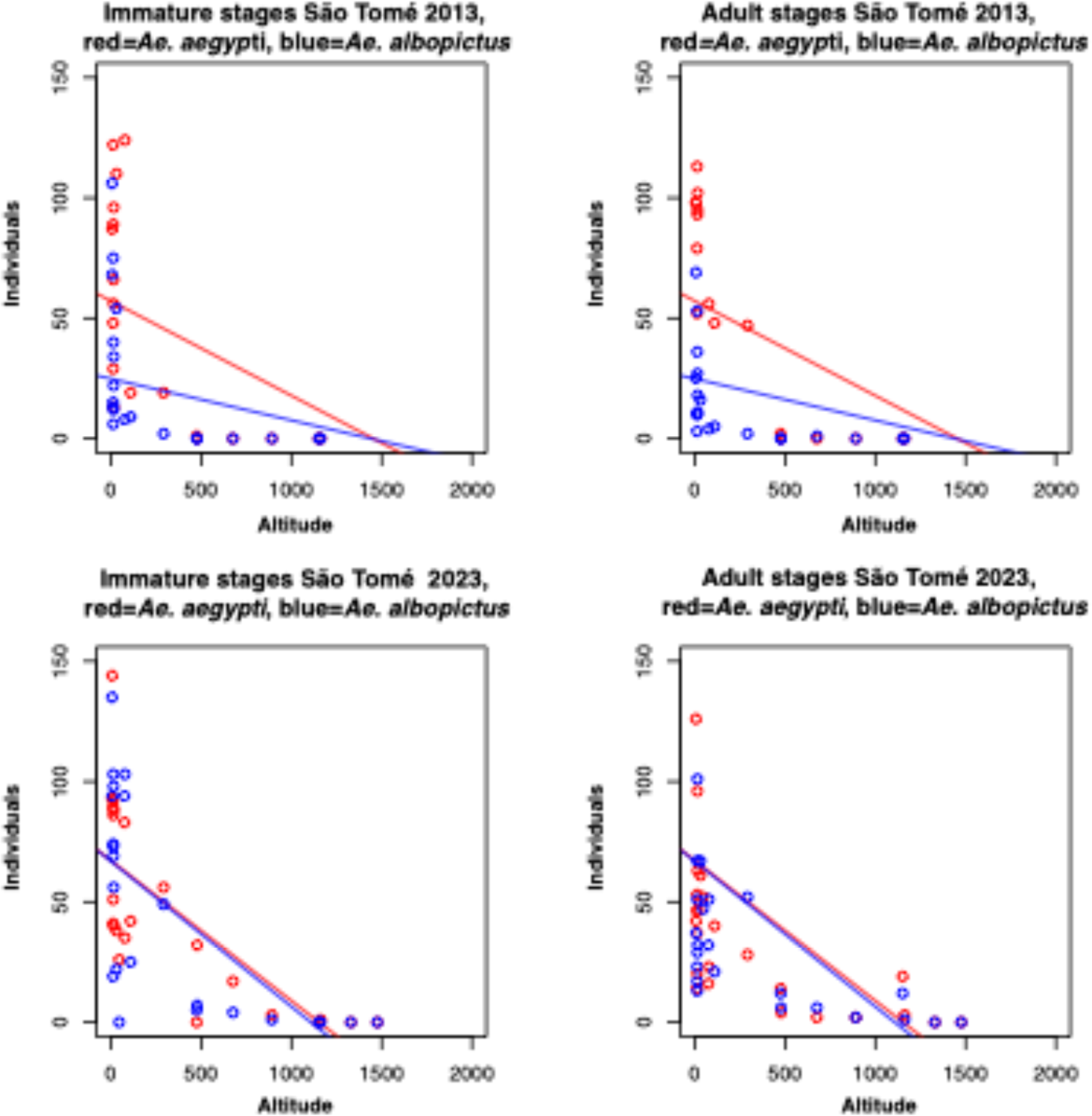
Abundance of two different species of *Aedes* along the altitudinal gradient in São Tomé island in two different years. **A.** Immature stages in 2013. **B.** Adults in 2013. **C.** Immature stages in 2023. **D.** Adults in 2023. Red: *Ae. aegypti*. Blue: *Ae. albopictus*. Lines represent the best fitting linear regression for each dataset.

We expanded the collection scheme from 2013 to a repeated sampling across the island. This allowed us to generate a crude comparison of the proportional representation of this vector across different locations in the island. Notably, we found a larger number of total specimens in 2013 (*n*_immatures_ = 1,557; *n*_adults_ = 3,308) than in 2023 (*n*_immatures_ = 2,088; *n*_adults_=1,666). Note that these collections do not take into account the potential for within-year variation and a fully-longitudinal study is needed to determine the potential disease burden and rate of progression of *Ae. albopictus* in the island of São Tomé. Using these counts, we did two analyses. First, we determined whether the abundance of each of the two *Aedes* species was affected by altitude. Our expectation was that the abundance of the vector would show an inverse relationship with altitude as has been observed in other locations (e.g. (Romiti *et al*. 2022)). Indeed, we find that in a linear model, the effect of altitude is significant for both life stages for the two sampled years (Table 1). The effect of species is significant in 2013 but not in 2023 which is consistent with the expansion of *Ae. albopictus* along the altitudinal gradient in the last decade. Contrary to our expectations that altitude would affect the abundance of the two species differently, we find no strong effect of the species × altitude interaction.

**TABLE 1.** Results of two linear models, one per year, describing the effect of altitude and species identity in abundance across the altitudinal gradient of São Tomé.

| Stage | Year | Altitude | Species | Species × altitude |
| --- | --- | --- | --- | --- |
| Immature | 2013 | $F=16.887, P = 2.188 \times 10^{-4}$ | $F = 6.663, P = 0.014$ | $F = 2.522, P = 0.121$ |
| Adult | 2013 | $F = 4.4857, P = 0.041$ | $F = 6.213, P = 0.017$ | $F = 2.545, P = 0.119$ |
| Immature | 2023 | $F = 39.906, 1.389 \times 10^{-7}$ | $F = 0.016, P = 0.902$ | $F = 0.013, P = 0.911$ |
| Adult | 2023 | $F = 20.413, 4.65 \times 10^{-5}$ | $F = 0.056, P = 0.814$ | $F = 0.029, P = 0.865$ |

Second, we studied whether the proportional representation of *Ae. albopictus* has changed over the years. We focused on the change that has occurred in the decade of sampling and analyzed the data of 2013 and 2023. First, we analyzed the data per year, pooling all traps. Pooled data suggested that in total the representation of *Ae. albopictus* has increased over time for both larvae (*Ae. albopictus*_2013_ = 29.80%, *Ae. albopictus*_2023_ = 49.378; 2-sample test for equality of proportions with continuity correction: χ^2^ = 140.48, df = 1, *p* < 1 × 10^-10^), and adults (*Ae. albopictus*_2013_ = 8.77%, *Ae. albopictus*_2023_ = 51.50%; 2-sample test for equality of proportions with continuity correction: χ^2^= 1,137.3, df = 1, *p* < 1 × 10^-10^). Second, we assess whether there had been a change in the relative composition at different locations along the altitudinal gradient. The proportional representation of *Ae. albopictus* for each location did not change measurably at the local scale as measured by the presence of immature stages (*t* = - 1.061, df = 12, *p* = 0.845) but did so at the level of adult individuals (*t* = 2.7938, df = 14, *p* = 7.177 × 10^-3^; Figure 3). These results suggest that the proportion of *Ae. albopictus* has increased in the last decade across the whole island, but that the vector community dynamics is largely determined at the local level; systematic samplings across the whole range are warranted.

**FIGURE 3.**
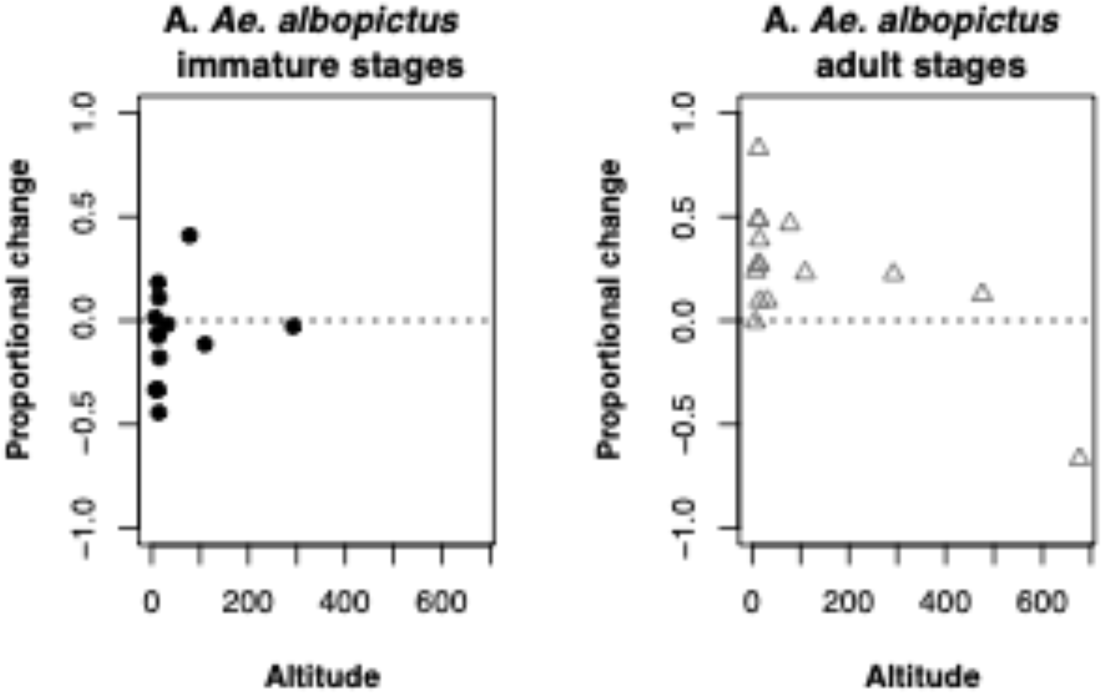
Change of *Ae.albopictus* over the 2013-2023 The y axis represents the difference in the proportional representation of *Ae. albopictus* individuals vs a second species, *Ae. aegypti* between 2023 and 2013. Positive values indicate an increase in 2023. **A.** Larval collections (*n* = 13), Adult collections (*n* = 15).The dashed line represents no change in the *Ae. albopictus* proportion.

A sampling on the island of Príncipe also revealed the presence of *Ae. albopictus* in Santo Antonio, the largest human settlement in Príncipe !“#”$%&’(#) +,-.//0. This samplingwas limited to adults only and we can’t confirm the presence of larval stages in the island. Even though this sampling is limited, as it only pertains to adults within a single year, it revealed the expansion of *Ae. albopictus* geographic range.

We used mitochondrial barcoding to provide a general view of the genealogical relationships between the isolates in São Tomé. The resulting tree was deposited in Open Tree of Life (Accession number TBD). We adhere to the haplotype nomenclature proposed in (Toto *et al*. 2003). We found individuals from the three mitochondrial haplotypes in São Tomé. Consistent with previous observations there is some geographical affinity for different haplotypes: ST1 is mostly found in the West slopes of Pico São Tomé, and ST3 is mostly found in the East side of the island (Table 2). We cannot determine whether this suggestive differentiation is related to geography or to the density of human settlements as the majority of human populations are located in the East side. These three haplotypes are all closely related and they form a monophyletic group along with a haplotype sampled in the Democratic Republic of Congo. Sampling of the island of Príncipe revealed the presence of haplotype ST1 but also of two more haplotypes not found in São Tomé and more closely related to a Republic of Congo haplotype than to Saotomean haplotypes (Figure 4). These results suggest the possibility of an independent colonization of the islands of São Tomé and Príncipe, and the possibility of a double colonization of Príncipe. Note that the level of variation in the COI locus is low and the branch-support is consequently limited. Conclusive studies about the demographic events and patterns of colonization of the islands of the Gulf of Guinea will require a more systematic sampling along the genome.

**FIGURE 4.**
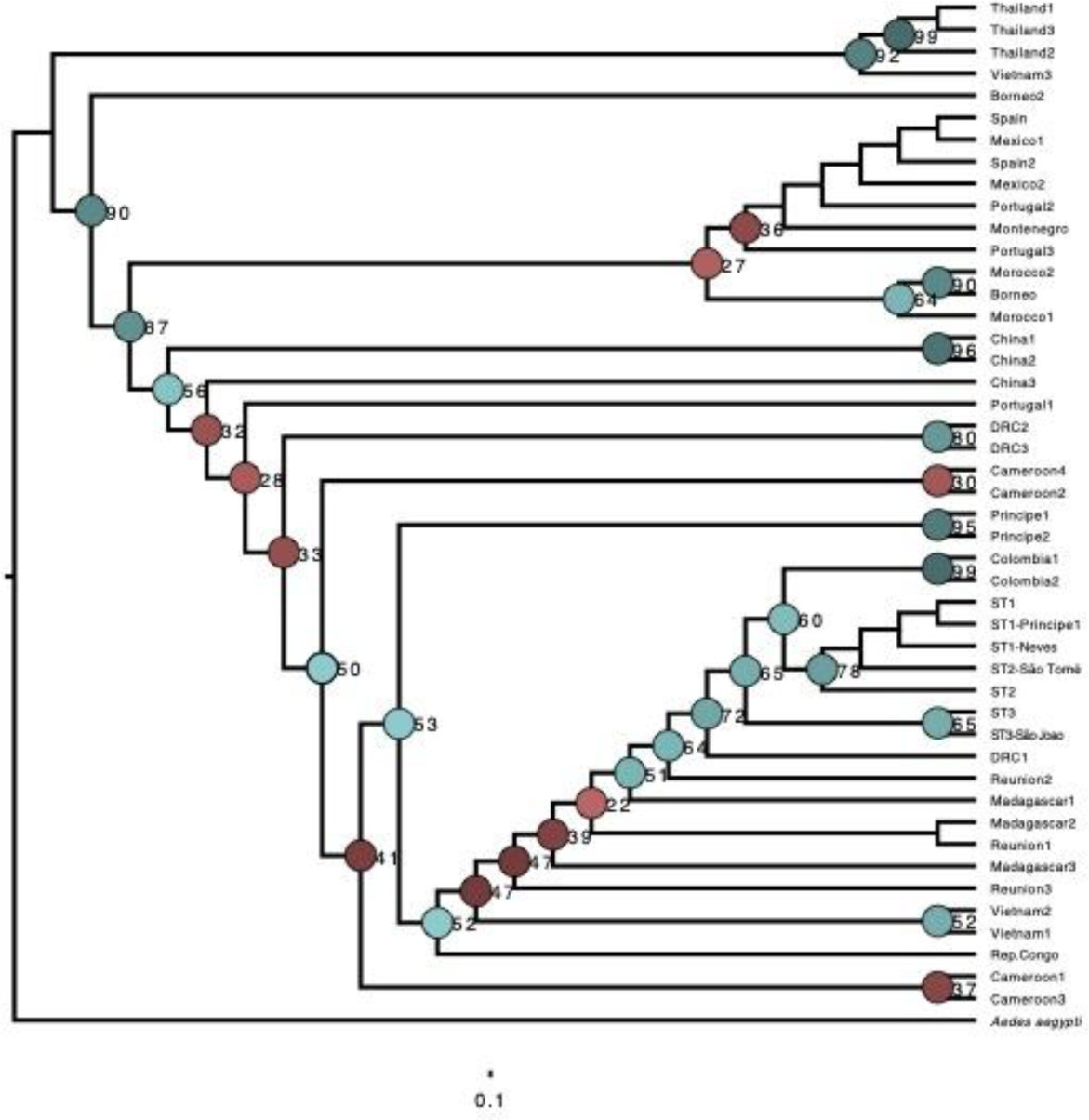
Maximum likelihood phylogenetic tree based on a COI barcode. Numbers above the nodes represent the ultrafast bootstrap branch support (1,000 replicates). Branches with bootstrap support higher than 50% are labeled with blue circles; labels with bootstrap support lower than 50% are labeled with red circles. GenBank accession numbers are listed in Table S1. ST: São Tomé, Rep. Congo: Republic of Congo, DRC: Democratic Republic of Congo.

**TABLE 2.** Haplotype frequency in *Ae. albopictus* from five locations in São Tomé and one location in Príncipe.

| Haplotype | NCBI accession number | Location |  |  |  |  |  |
| --- | --- | --- | --- | --- | --- | --- | --- |
|  |  | São Tomé | Neves | São Joao dos Angolares | Santo Amaro | Trindade | Principe |
| ST1 | JF309317 | 2 | 22 | 5 | 5 | 11 | 2 |
| ST2 | JF309319 | 9 | 8 | 16 | 14 | 9 | 0 |
| ST3 | JF309318 | 19 | 6 | 10 | 9 | 17 | 0 |
| Principe1 | TBD | 0 | 0 | 0 | 0 | 0 | 7 |
| Principe2 | TBD | 0 | 0 | 0 | 0 | 0 | 6 |
| *n* |  | 30 | 36 | 31 | 28 | 37 | 15 |

Next we inferred habitat suitability, and thus the potential ranges for *Aedes* on São Tomé and Príncipe using a maximum entropy analysis. Our MaxEnt model, based on 79,247 global observations of *Ae. albopictus*, yielded a global prediction of habitat suitability (see Supplemental Figure S1). The MaxEnt model included 19,595 training globally distributed samples and 28,223 background points over 500 iterations. The training AUC was 63.5%. We find that the climate on São Tomé and Príncipe is broadly suitable for *Aedes* to persist and expand its range across the islands. Though our sampling efforts did not find mosquitoes above approximately 1350 m of elevation, the suitable bioclimatic range indicated by the MaxEnt model extends to the top of Pico São Tomé, the highest point on the island. The maximum elevation on Príncipe is much lower than on São Tomé, and thus less likely to limit the range of *Ae. albopictus*.

Our global MaxEnt model identified five climatic variables which explained a cumulative 87% of the overall variance (Figure 5 and Table S2) that predict the presence of *Ae. albopictus*, including mean temperature of the coldest quarter (Bio11) and four variables describing precipitation: precipitation of the coldest quarter (Bio19), annual precipitation (Bio12), precipitation of the driest quarter (Bio17), and precipitation of the driest month (Bio14). In all cases, the range of values that *Aedes* are exposed to on the islands is smaller than the species experiences around the world, and in three of these variables (precipitation of the coldest quarter, precipitation of the driest quarter, and precipitation of the driest month) the range of values on the islands is completely encompassed within the global range (Figure 6). Mean temperature of the coldest quarter and annual precipitation on the islands overlap strongly with, but extend beyond the upper thresholds of the global ranges (Figure 6). Taken together, these results indicate that the *Ae. albopictus* populations on São Tomé and Príncipe are unlikely to face the same thermal or precipitation limitations that appear to constrain global populations. Our samples on São Tomé capture at least 63.7% for each of the five bioclimatic axes, and the model results suggest that *Aedes* could spread beyond the sampled geographic range if it has not already done so.

**FIGURE 5.**
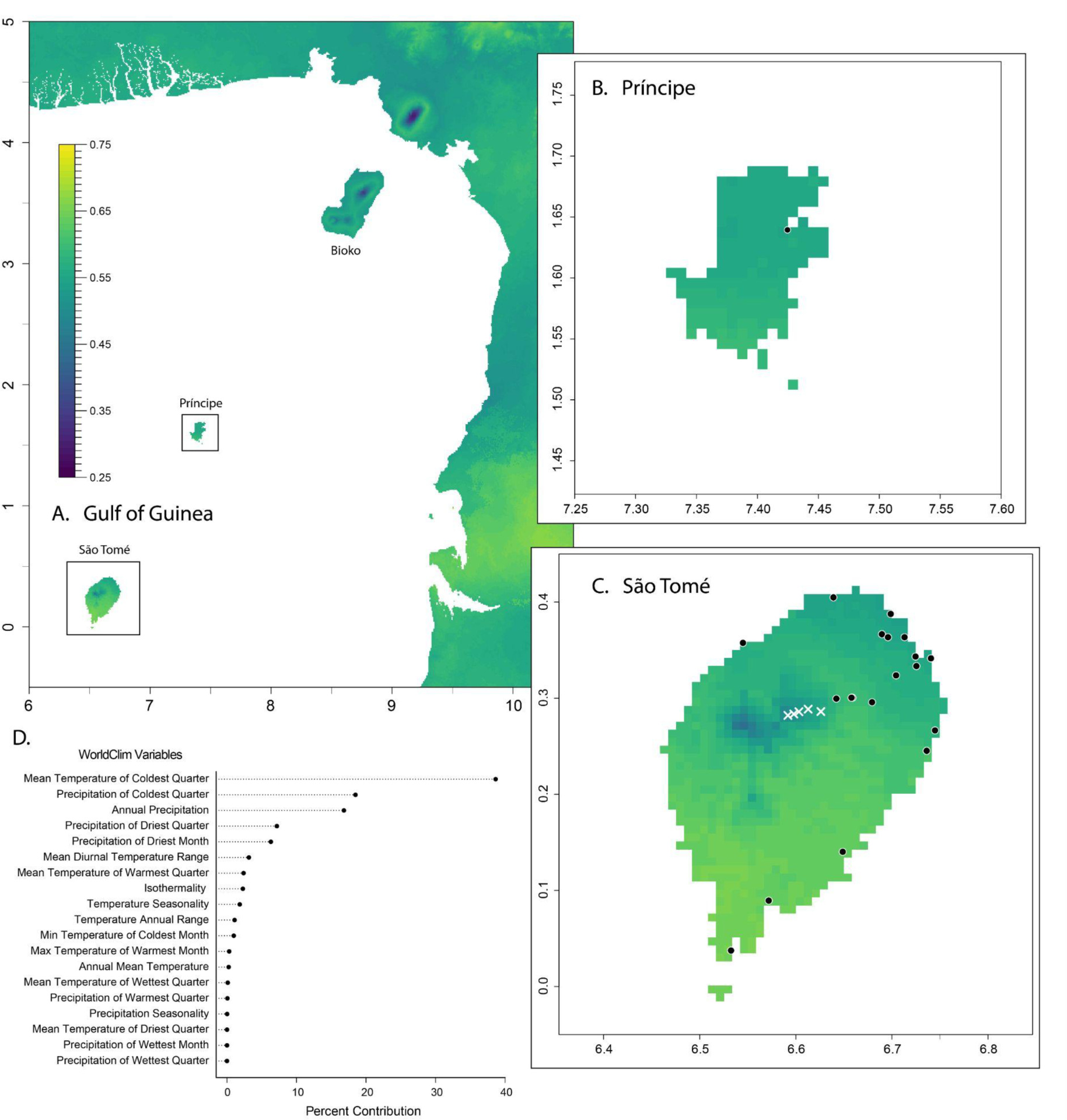
Map depicting the results of a maximum entropy (MaxEnt) model for the Gulf of Guinea (A) and the islands of Príncipe (B) and São Tomé (C). Heatmap coloration shows predicted habitat suitability for *Aedes albopictus* mosquitoes based on bioclimatic data from the WorldClim database, with bluer colors representing relatively poor habitat and the yellow end of the gradient depicting more suitable habitat. Data points in panes B and C show presence (black data points) or absence (white crosses) of *Ae. albopictus* from our sampling efforts. The contribution of each WorldClim variable to predicting favorable *Ae. albopictus* habitat is shown in pane D.

**FIGURE 6.**
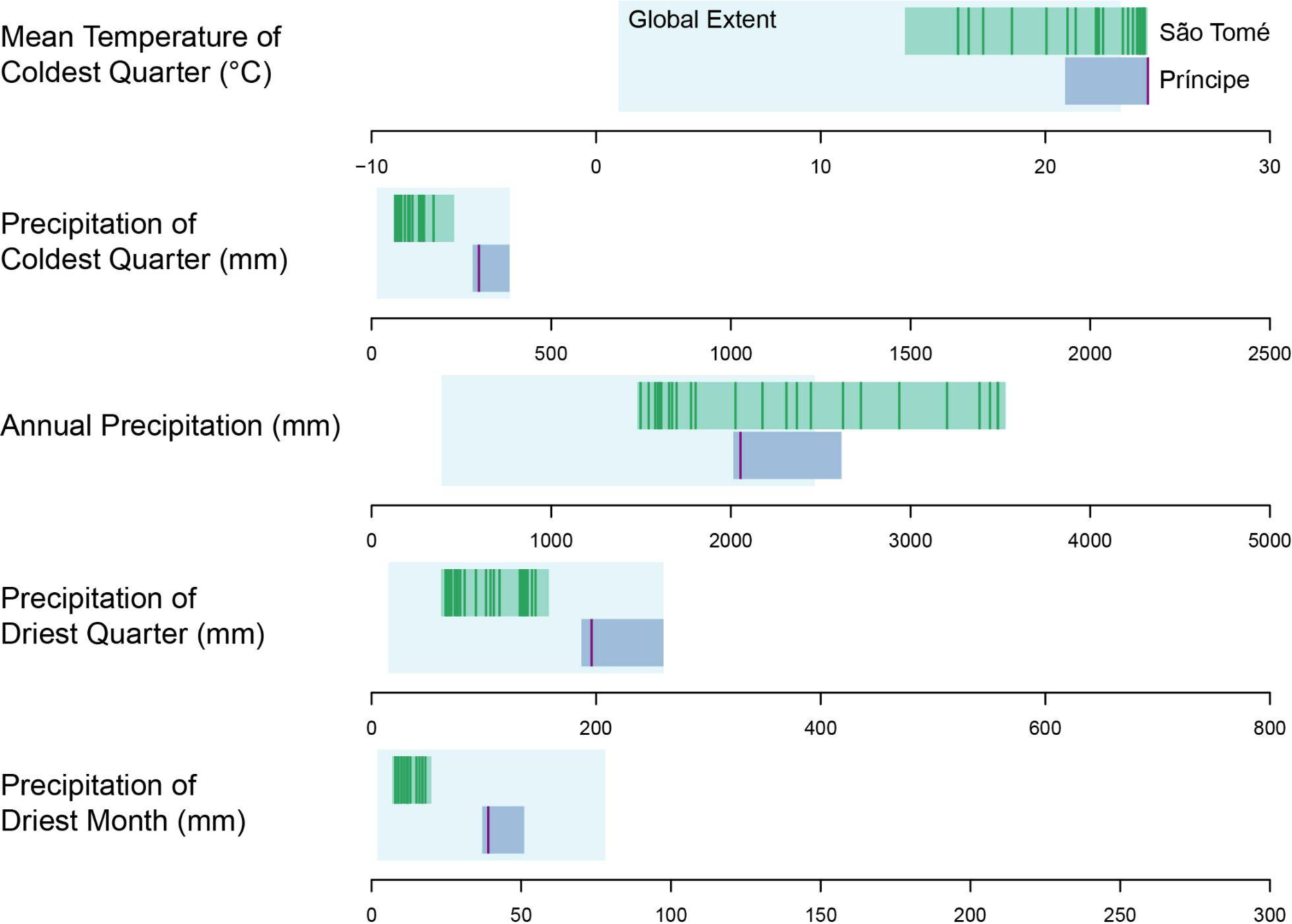
Bioclimatic niche axes for *Aedes albopictus* on São Tomé and Príncipe. MaxEnt modeling identified five bioclimatic niche axes that are associated with global suitability of habitat for *Ae. albopictus* (see Figure 5). The pale blue boxes depict the central 95% quantile range (discarding the upper and lower 2.5%) of the global distribution of *Ae. albopictus* on each bioclimatic axis. The extent of each of the axes is represented with a green box for São Tomé and with a blue box for Príncipe. Vertical hash marks within each island box show the values at each of our sampling localities.

## DISCUSSION

The range of disease vectors is critical to determine the disease risk of different human populations. The last systematic assessment of the vectors in the island of São Tomé reported the presence of rare *Aedes* species endemic to the Gulf of Guinea (Mourão 1964; Loiseau *et al*. 2022) but not of *Ae. albopictus*. Our study follows on a previous report that established the presence of *Ae. albopictus* in the island of São Tomé (Reis *et al*. 2017). We expanded the previous sampling, and sampled across an altitudinal gradient to determine the current and potential future range of this vector species in São Tomé. We found that the potential for spread is high and might encompass high elevation regions. We captured *Ae. albopictus* in many of the main population centers of the island, São Tomé (“#”$%&’(#) + 53,000), Santo Amaro (8,300), Neves (7,500), Santana (7,000), and Trindade (6,600) but also in mid-elevation villages (e.g., Monte Cafe, Trindade), which might suggest the high prevalence of the mosquito across São Tomé.

A novel observation is the presence of *Ae. albopictus* in a third island of the Gulf of Guinea, Príncipe. Besides São Tomé, previous reports have also found *Ae. albopictus* on the island of Bioko (Toto *et al*. 2003), a land-bridge island in Equatorial Guinea that is ∼30km from the coast of Cameroon in 2001. Given the proximity of Bioko to the mainland, it seems possible that the colonization of the islands in the Gulf of Guinea has followed a stepping-stone model of colonization. Nonetheless, the sparsity of the collections reported so far do not allow us to test this possibility. A natural followup question is whether *Ae. albopictus* has already colonized the island of Annobón, the fourth and most remote of the islands in the Cameroon Volcanic Line.

*Aedes albopictus* has been successful at the colonization of almost every single country in Africa (Longbottom *et al*. 2023). The first reports on the African continent date from the early 1990s in Nigeria (Savage *et al*. 1992) and the firsts reports in countries of the Gulf of Guinea came a few years later (Cameroon 1999, (Fontenille and Toto 2001); Equatorial Guinea, (Toto *et al*. 2003)). The species has also colonized several of the African islands. The species was first described in Mayotte in 2007, Beilhe et al., 2012 and in Réunion Island, (Delatte et al., 2008). Notably the species has been present in the Seychelles and Mauritius for longer (Rai 1986) and in Madagascar for even longer (Fontenille and Rodhain 1989). Ecologically, these locations are diverse (e.g, latitude differences span almost the totality of the Southern Hemisphere). The large variety of environments in which *Ae. albopictus* has been found underscores the ability of this species to adapt to novel habitats.

Environmental niche modeling indicates that the climate on both São Tomé and Principé is broadly suitable to sustaining populations of *Ae. albopictus*. The best predictors of global habitat suitability in our MaxEnt model appeared to center on the minimum temperature and precipitation that the mosquitoes would have to endure through the year. Though these factors are limiting globally, neither São Tomé nor Príncipe display values near the low end of the global ranges occupied by *Ae. albopictus* mosquitoes. Indeed, the climate on the islands offers greater mean annual temperature, and more precipitation than *Ae. albopictus* experience elsewhere. Therefore, climate is unlikely to limit the spread of this vector on São Tomé or Príncipe. Our prediction of the potential spread of *Ae. albopictus* is not without caveats. The bioclimatic model did not account for other niche factors such as habitat type, anthropogenic land use, availability of host animals, or human commensalism in the mosquitoes, and these potential constraints on the spread of *Ae. albopictus* merit further attention. Though it may not capture the totality of important niche characteristics, ecological niche modeling like we present here is an important tool in understanding the human disease risk posed by insect-vectored diseases.

The implications of finding *Ae. albopictus* in the island of Príncipe and mid-elevations in São Tomé is of importance for public health. Dengue presented itself as an outbreak in the island of São Tomé in 2021 (Carvalho *et al*. 2022). In 2022, there were 1,161 cases of Dengue diagnoses in the island (Lessa *et al*. 2023). The disease remains woefully undiagnosed in the rest of the archipelago. Moreover, *Ae. albopictus* is a competent vector of Zika and Chikunguya (Gloria-Soria *et al*. 2020) and has been implicated in outbreaks of Chikunguya in countries of Central Africa (e.g., Gabón, Cameroon, (Demanou *et al*. 2010; Paupy *et al*. 2012; Grard *et al*. 2014; Ngoagouni *et al*. 2015; Obame-Nkoghe *et al*. 2023)). While there have been no cases of Zika or Chikunguya reported for São Tomé and Príncipe, there is a new potential vector for these diseases in the country and surveillance policies should include training to identify this species and screen for relevant infections. An additional, but related aspect of these assessments would also include the extent of genetic variability within each species. In the case of vectors, genetic variability might translate into insecticide resistance, ability to colonize even further habitats, and higher attraction to humans.

Ecological studies have suggested that the colonization of *Ae. albopictus* can have important epidemiological consequences. *Aedes albopictus*, a species historically considered to be a secondary and less effective vector of dengue and yellow fever (reviewed in (Gratz 2004)), might displace *Ae. aegypti* (the primary vector of these diseases) in places where they concur. In Seychelles, it has been hypothesized that *Ae. albopictus* has excluded *Ae. aegypt*i from the granitic islands (Le Goff *et al*. 2012). A similar displacement has been proposed for the island of Mayotte (Bagny *et al*. 2009). Since *Ae. albopictus* has proven to be an effective vector of dengue and other diseases (Jupp and Kemp 2002; Garcia-Luna *et al*. 2018; McKenzie *et al*. 2019; Gloria-Soria *et al*. 2020), colonizations of a new range by *Ae. albopictus* are likely to not be benign. Only systematic studies of disease burden coupled with longitudinal studies of the presence of different vectors will be able to assess the risk of the resident population of São Tomé and Príncipe.

Our counts are in line with previous studies in other locations that show that *Aedes* species are highly restricted to low and mid elevations. The highest elevation at which *Aedes* was collected in São Tomé was 677 m above sea level. Nonetheless, collections of *Aedes* at high altitude have been reported in Bolivia (Ríos *et al*. 2023), Kenya (Hertz *et al*. 2016), and Colombia (Ruiz-López *et al*. 2016). As climate change progresses and both *Ae. aegypti* and *Ae. albopictus* increase their population sizes, which in turn gives the chance for further adaptation, it is not unlikely that these vector species might expand to high altitude environments.

