## Supplementary material for "The spread of *Aedes albopictus* in the islands of São Tomé and Príncipe": Table S1, Figure S1

**FIGURE S1.** Global occurrence data for *Aedes albopictus* since 1980 from the Global Biodiversity Information Facility (GBIF; pane A). We used the occurrence data, paired with bioclimatic data to construct a global maximum entropy model (pane B) which we used to predict habitat suitability on São Tomé and Príncipe.

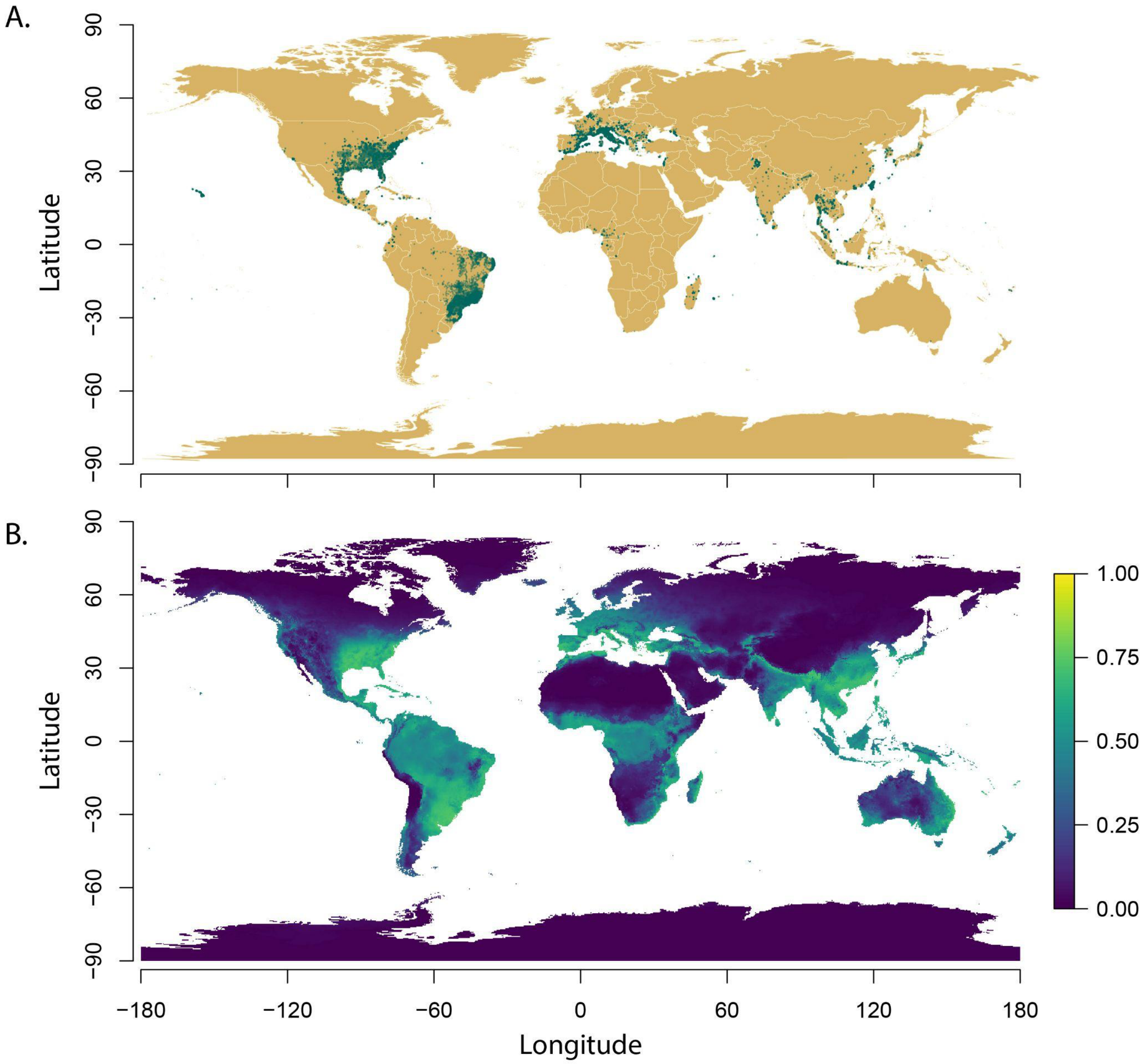
A.
B.
Latitude
90
60
30
0
−30
−60
−90
Longitude
−180
−120
−60
0
60
120
180
1.00
0.75
0.50
0.25
0.00

**TABLE S1.** GenBank accession numbers for the COI barcodes used to generate an *Ae. albopictus* gene genealogy.

| **Code in Figures 2 and S1** | **GenBank accession number** |
|---|---|
| Thailand1 | KM613129.1 |
| Thailand3 | KM613121.1 |
| Thailand2 | KM613128.1 |
| Vietnam3 | HQ398902.1 |
| Borneo2 | MN540322.1 |
| Spain | KU319448.1 |
| Mexico1 | MT999274.1 |
| Spain2 | KU319443.1 |
| Mexico2 | MT552470.1 |
| Portugal2 | MK995331.1 |
| Montenegro | MK505589.1 |
| Portugal3 | MK995330.1 |
| Morocco2 | KU522419.1 |
| Borneo | MN540323.1 |
| Morocco1 | KU522421.1 |
| China3 | KX886337.1 |
| China1 | KX981869.1 |
| China2 | KX981868.1 |
| Portugal1 | MK995332.1 |
| DRC2 | MT345388.1 |
| DRC3 | MT345383.1 |
| Cameroon4 | MH921571.1 |
| Cameroon2 | MH921570.1 |

| | |
|---|---|
| Principe1 | TBD |
| Principe2 | TBD |
| Colombia1 | KP877575.1 |
| Colombia2 | KP877572.1 |
| ST1 | JF309319.1 |
| ST2 | JF309319.1 |
| ST3 | JF309318.1 |
| DRC1 | MT345390.1 |
| Reunion2 | AJ971012.1 |
| Madagascar1 | AJ971007.1 |
| Madagascar2 | JN406732.1 |
| Madagascar3 | JN406725.1 |
| Reunion1 | JN406663.1 |
| Reunion3 | AJ971013.1 |
| Vietnam2 | KX573911.1 |
| Vietnam1 | KX495925.1 |
| Rep. Congo | MH025948.1 |
| Cameroon1 | MH921572.1 |
| Cameroon3 | MH921568.1 |
| *Aedes aegypti* | MK265729.1 |

**TABLE S2.** WorldClim variables included in MaxEnt model, and their contributions to the final model. Bolded text denotes variables with high contribution.

| WordClim Variable | Inclusion | Contribution |
|---|---|---|
| BIO1 = Annual Mean Temperature | Included | 0.2451 |
| BIO2 = Mean Diurnal Range (Mean of monthly (max temp - min temp)) | Included | 3.1556 |
| BIO3 = Isothermality (BIO2/BIO7) (×100) | Excluded | 2.2919 |
| BIO4 = Temperature Seasonality (standard deviation ×100) | Included | 1.8381 |
| BIO5 = Max Temperature of Warmest Month | Included | 0.3236 |
| BIO6 = Min Temperature of Coldest Month | Included | 0.9791 |
| BIO7 = Temperature Annual Range (BIO5-BIO6) | Excluded | 1.1003 |
| BIO8 = Mean Temperature of Wettest Quarter | Included | 0.1155 |
| BIO9 = Mean Temperature of Driest Quarter | Included | 0.0116 |
| BIO10 = Mean Temperature of Warmest Quarter | Included | 2.3871 |
| **BIO11 = Mean Temperature of Coldest Quarter** | **Included** | **38.6427** |
| **BIO12 = Annual Precipitation** | **Included** | **16.8210** |
| BIO13 = Precipitation of Wettest Month | Included | 0.0094 |
| **BIO14 = Precipitation of Driest Month** | **Included** | **6.3165** |
| BIO15 = Precipitation Seasonality (Coefficient of Variation) | Included | 0.0269 |
| BIO16 = Precipitation of Wettest Quarter | Included | 0.0000 |
| **BIO17 = Precipitation of Driest Quarter** | **Included** | **7.1773** |
| BIO18 = Precipitation of Warmest Quarter | Included | 0.0726 |
| **BIO19 = Precipitation of Coldest Quarter** | **Included** | **18.4860** |
